# Physiological reason for ceasing growth of unfertilized eggs produced by unmated queens in the subterranean termite *Reticulitermes chinensis*

**DOI:** 10.1101/037390

**Authors:** Ganghua Li, Long Liu, Pengdong Sun, Yao Wu, Chaoliang Lei, Xiongwen Chen, Qiuying Huang

**Affiliations:** College of Plant Science and Technology, Huazhong Agricultural University, Wuhan 430070, Hubei, China; College of Life Science, Hubei Normal University, Huangshi 435002, Hubei, China

**Keywords:** termites, asexual queen succession, ovarian development, embryonic growth, physiological indices

## Abstract

In *Reticulitermes chinensis,* a close relative of *R. speratus* with asexual queen succession, unfertilized eggs can be produced but are not incubated. To explain this phenomenon, we analysed the physiological differences between unfertilized eggs/unmated queens and fertilized eggs/mated queens. Fertilized eggs consumed significantly larger quantities of five amino acids (Cys, Met, Ile, Leu and Tyr), Ca, protein and cholesterol during incubation. The higher levels of four trace elements (Na, K, Zn and Fe) in fertilized eggs and their lower levels in mated queens indicated that mated queens might transfer these trace elements to fertilized eggs to complete incubation. The higher levels of Mn, triglycerides and serotonin in mated queens and higher levels of Mn and glucose in fertilized eggs suggested that these substances are very important for normal ovarian and embryonic growth. The different expression of three reproductive genes *(vtgl, rabil* and *JHE1)* suggested that they might be involved in the regulation of ovarian and embryonic growth. Overall, changes in these physiological indices may substantially affect ovarian and embryonic growth and prohibit the incubation of unfertilized eggs in *R. chinensis.*

## INTRODUCTION

Recently, asexual queen succession (AQS) has been described in three species of lower termites [*Reticulitermes speratus* (Matsuura et al., 2009), *R. virginicus* (Vargo et al., 2012) and *R. lucifugus* (Luchetti et al., 2013)] and two species of higher termites [*Embiratermes neotenicus* (Fougeyrollas et al., 2015) and *Cavitermes tuberosus* (Roisin et al., 2014)]. In AQS species of termites, workers, soldiers and alates are sexually produced but neotenic queens arise through thelytokous parthenogenesis (Matsuura et al., 2009). This AQS system enables the primary queen to maintain her full genetic contribution to the next generation while avoiding any loss of genetic diversity from inbreeding (Matsuura, 2011). Most of the work conducted on the AQS system in termites has focused on the discovery of new AQS species and the mechanism of facultative parthenogenesis (Fougeyrollas et al., 2015; Kawatsu and Matsuura, 2013; Luchetti et al., 2013; Roisin et al., 2014; Vargo et al., 2012; Yashiro and Matsuura, 2014). Some termite species that are evolutionally related to AQS species of termites have been demonstrated to exhibit neither AQS nor parthenogenesis (Kawatsu and Matsuura, 2013; Luchetti et al., 2013). However, little is known about why these termite species have no AQS and specifically why unfertilized eggs produced by unmated queens of these species are unable to hatch

The ovarian and embryonic development of insects are both complex physiological processes that are influenced by many physiological factors, including nutrients (Bodnaryk and Morrison, 1966; Judd et al., 2010), trace elements (Chaudhury et al., 2000; Hammack, 1999), hormones (Hagedorn et al., 1975; Lagueux et al., 1977) and reproductive genes (e.g., vitellogenin genes) (Ishitani and Maekawa, 2010; Maekawa et al., 2010). For example, wasp eggs require substantial quantities of proteins and lipids to provide the materials and energy for embryonic growth (Judd et al., 2010). Adult female house flies require an adequate protein diet to initiate and sustain normal ovarian development (Bodnaryk and Morrison, 1966). In adult mosquitoes (Hagedorn et al., 1975) and adult females of *Locusta migratoria* (Lagueux et al., 1977), ecdysone plays an important role in stimulating egg development and vitellogenin synthesis. Metal ions participate in the process of enzyme activation and in trigger and control mechanisms (Chaudhury et al., 2000). For example, potassium (K) was reported to be a key ion for protein-dependent egg maturation in three insects, *Phormia regina*, *Sarcophaga bullata*, and *Cochliomyia hominivorax* (Chaudhury et al., 2000; Hammack, 1999). Previous studies on ovarian development in termites have primarily focused on changes in ovarian morphology, JH titres and the expression of vitellogenin genes at different queen stages in *R. speratus* (Ishitani and Maekawa, 2010; Maekawa et al., 2010). With regard to embryonic growth in termites, differences in size, survival rate, and the length of the hatching period between unfertilized and fertilized eggs have been investigated in *R. speratus* (Matsuura and Kobayashi, 2007). However, the physiological differences between unfertilized eggs/unmated queens and fertilized eggs/mated queens, such as the content of amino acids, nutrients, trace elements, hormones, neurotransmitters and the expression of reproductive genes, have not been investigated in termites.

The subterranean termite *R. chinensis*, which is widely distributed in China, causes serious damage to buildings and forests and results in major economic losses (Liu et al., 2015). In the present study, we found that unfertilized eggs can be produced by unmated queens of *R. chinensis*, but they do not hatch under laboratory and simulated field conditions, suggesting that *R. chinensis* exhibits neither AQS nor parthenogenesis. To explore why unfertilized eggs of *R*. *chinensis* are unable to hatch, we conducted a comprehensive analysis of physiological differences in ovarian and embryonic growth between unfertilized eggs/unmated queens and fertilized eggs/mated queens. We found that unfertilized eggs ceased embryonic growth and had significant differences in morphological characters, size and micropyle (sperm gate) number compared with fertilized eggs in the final stage (stage V) of development. Moreover, there were significant differences in the levels of 11 amino acids, six trace elements, four nutrients, serotonin, and the expression of three reproductive genes (*vtg 1*, *rab 11 and JHE1*) between unfertilized eggs/unmated queens and fertilized eggs/mated queens, suggesting that the absence of incubation of unfertilized eggs produced by unmated queens is associated with the above changes in physiological indices in *R. chinensis*. Our physiological findings contribute to an understanding of why the subterranean termite *R. chinensis* exhibits neither AQS nor parthenogenesis even though it is a close relative of *R. speratus*, which does exhibit AQS (Austin et al., 2004; Matsuura et al., 2009).

## RESULTS

### Formation of female-female colonies and female-male colonies under laboratory and simulated field conditions

The number of unfertilized eggs in FF colonies was significantly higher than the number of fertilized eggs in FM colonies at stages I and II (Fig. 1A; stage I: *p* = 0.001; stage II: *p* < 0.001). The numbers of both unfertilized and fertilized eggs decreased during stages III and IV, and no significant differences were found (Fig. 1A; stage III: *p* = 0.643; stage IV: *p* = 0.214). At stage V, no newly produced eggs were found in either the FF or FM colonies.

**Fig 1:**
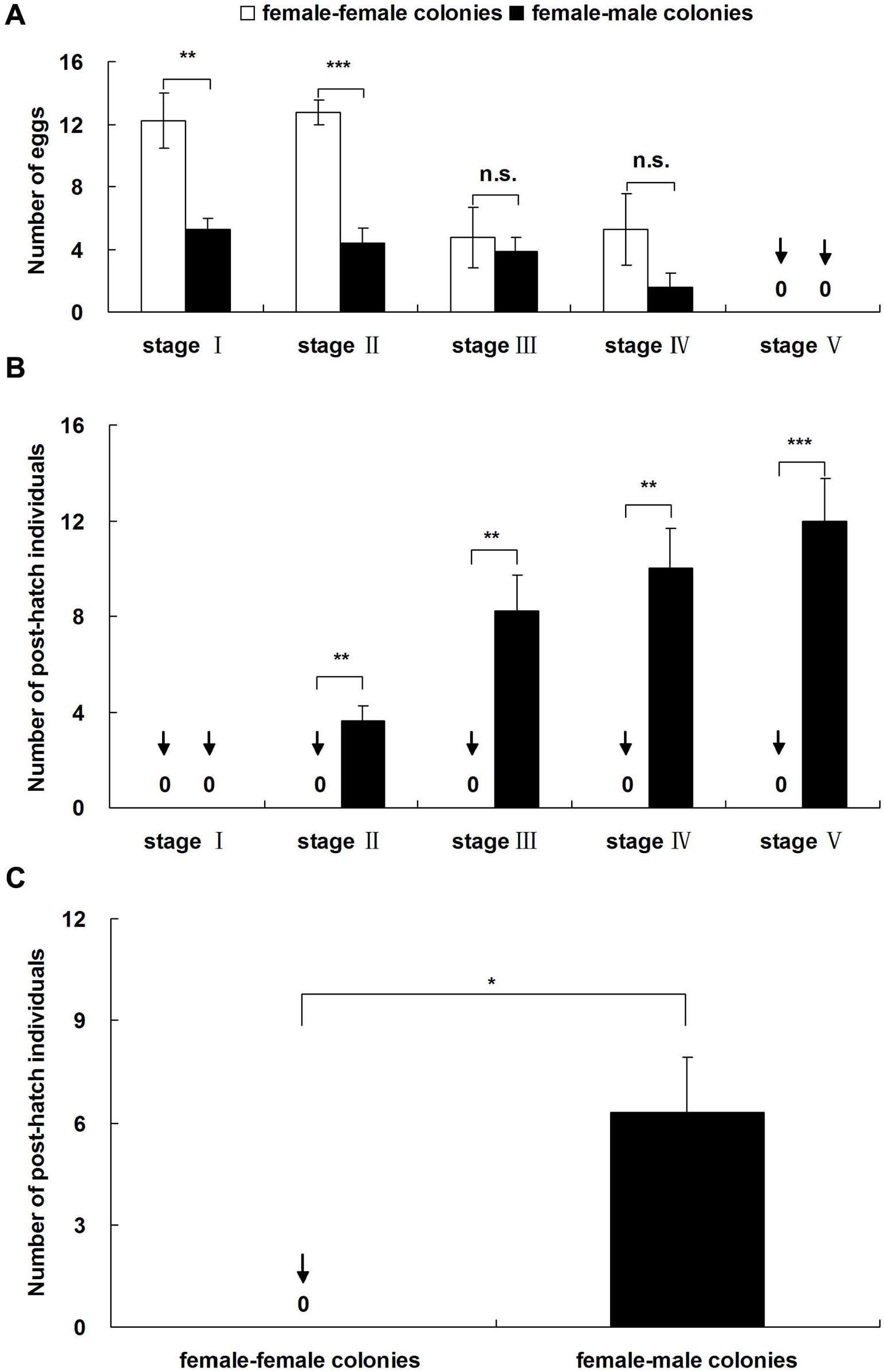
Comparison of brood number (mean ± S.E.M.) between female-female and femalemale colonies. (a) Eggs and (b) post-hatch individuals under laboratory conditions; (c) post-hatch individuals under simulated field conditions. Asterisks denote significant differences (paired *t*-test, n.s. = not significant, * *p* < 0.05, ** *p* < 0.01, *** *p* < 0.001).

No post-hatch individuals were observed in any FF colony at all five developmental stages (Fig. 1B). Post-hatch individuals (larvae, workers, pre-soldiers or soldiers) were found after stage II in the FM colonies (Fig. 1B). Larvae (1-5 individuals per colony) were found at stage II in all eight FM colonies (8/8 colonies: 100%). At stage III, all the eight FM colonies (100%) contained workers (2-8 individuals per colony). Two pre-soldiers were found in two of the FM colonies at stage IV (2/8 colonies: 25%). Five soldiers were found in five colonies during stage V (5/8 colonies: 62.5%). The number of post-hatch individuals in the FM colonies was significantly higher than in FF colonies for stages II to V (Fig. 1B; stages II-V: *p* < 0.01).

In simulated field conditions, no post-hatch individuals were found in any of the FF colonies 5.5 months after colony formation. However, post-hatch individuals (average 6.29 individuals per colony) were found 5.5 months after colony formation in all the FM colonies (Fig. 1C). The number of post-hatch individuals in the FM colonies was significantly higher than in the FF colonies (Fig. 1C; *p* = 0.024).

### Morphological observation of eggs at the five developmental stages

The embryos of fertilized eggs developed normally and exhibited visible differences among the five developmental stages (from stage I to stage V, Fig. 2A). In unfertilized eggs, most of embryos ceased development at stage II, and only a few of the embryos developed to stage III or stage I V, but none of the embryos developed to stage V, unlike the fertilized eggs (Fig. 2A). There were no significant differences in morphological characters and size between unfertilized eggs and fertilized eggs prior to stage IV (Fig. 2A and B). However, the unfertilized eggs gradually shrank and spoiled, and they were significantly smaller than fertilized eggs at stage V (Fig. 2A and C; *p* = 0.008). Micropyles were located on the posterior end of the eggs. Almost all the unfertilized eggs (14/15) had no micropyles, but all fertilized eggs had micropyles (Fig. 2D).

**Fig 2:**
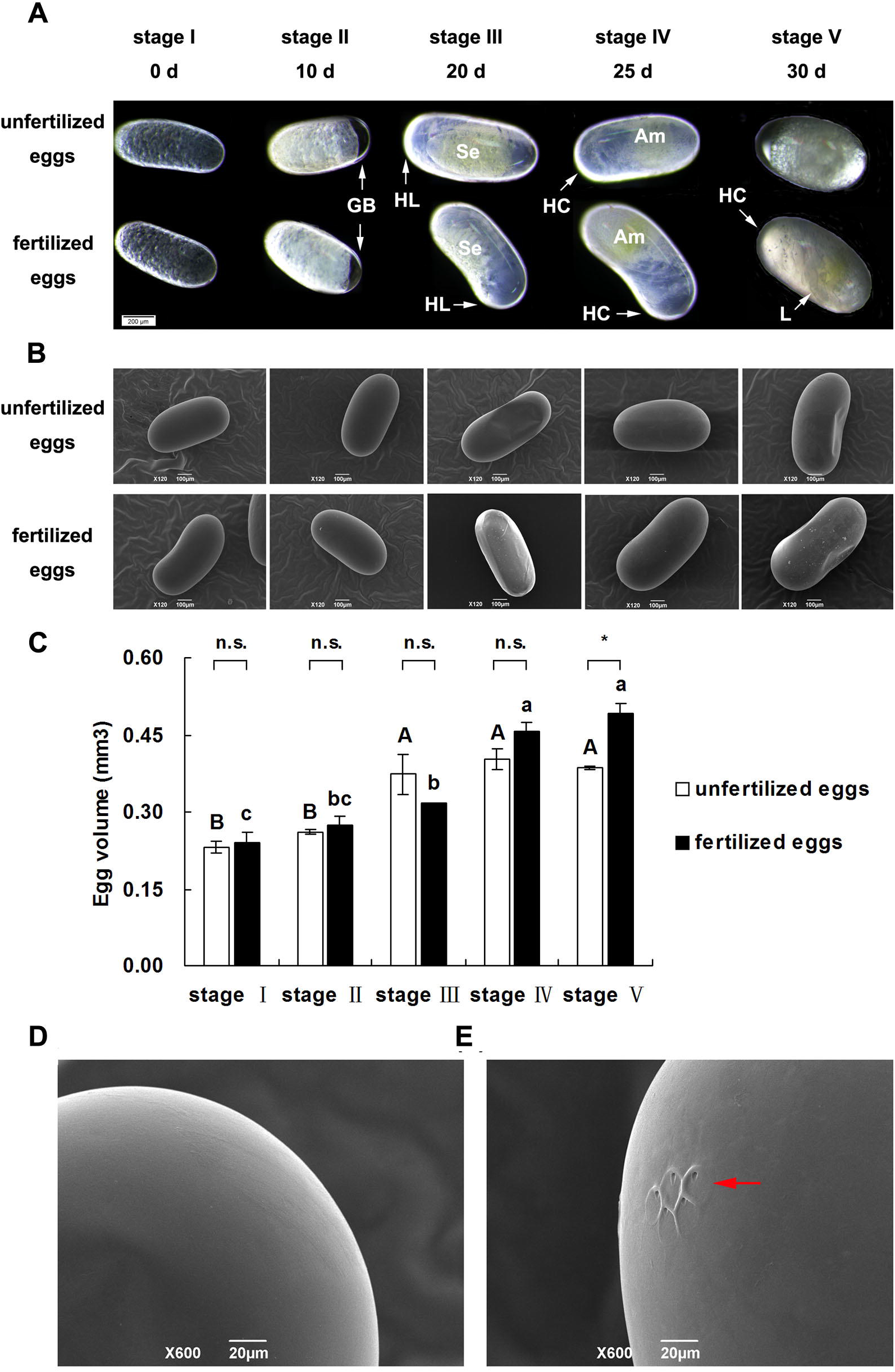
Differences in morphological characters, size and micropyle number between unfertilized and fertilized eggs. (a) Egg morphology under stereoscope. GB: germ band; HL: head lobe; HC: head capsule; Se: serosa; Am: amnion; L: leg; (b) egg morphology under SEM; (c) differences in egg size between unfertilized and fertilized eggs at five developmental stages under SEM; (d) unfertilized eggs without micropyles; and (e) fertilized eggs with micropyles under SEM.

### Amino acids

The levels of five amino acids (Cys, Met, Ile, Leu and Tyr) in unfertilized eggs from FF colonies were significantly higher than in fertilized eggs from FM colonies (Fig. 3A and Table S1; Cys: *t* = -8.000, *df* = 2, *p* = 0.015; Met: *t* = -6.424, *df* = 2, *p* = 0.023; Ile: *t* = -10.250, *df* = 2, *p* = 0.009; Leu: *t* = -5.277, *df* = 2, *p* = 0.034; Tyr: *t* = -9.430, *df* = 2, *p* = 0.011). The levels of two amino acids (Asp and Ala) in unmated queens were significantly lower than in mated queens (Fig. 3B; Asp: *t* = 4.014, *df* = 2, *p* = 0.016; Ala: *t* = 10.123, *df* = 2, *p* = 0.001), but the levels of six amino acids (Gly, Ile, Leu, Phe, His and Arg) in unmated queens were significantly higher than in mated queens (Fig. 3B; Gly: *t* = -5.354, *df* = 2, *p* = 0.006; Ile: *t* = -15.296, *df* = 2, *p* < 0.001; Leu: *t* = -3.101, *df* = 2, *p* = 0.036; Phe: *t* = -4.177, *df* = 2, *p* = 0.014; His: *t* = -5.494, *df* = 2, *p* = 0.005; Arg: *t* = -5.087, *df* = 2, *p* = 0.007).

**Fig 3:**
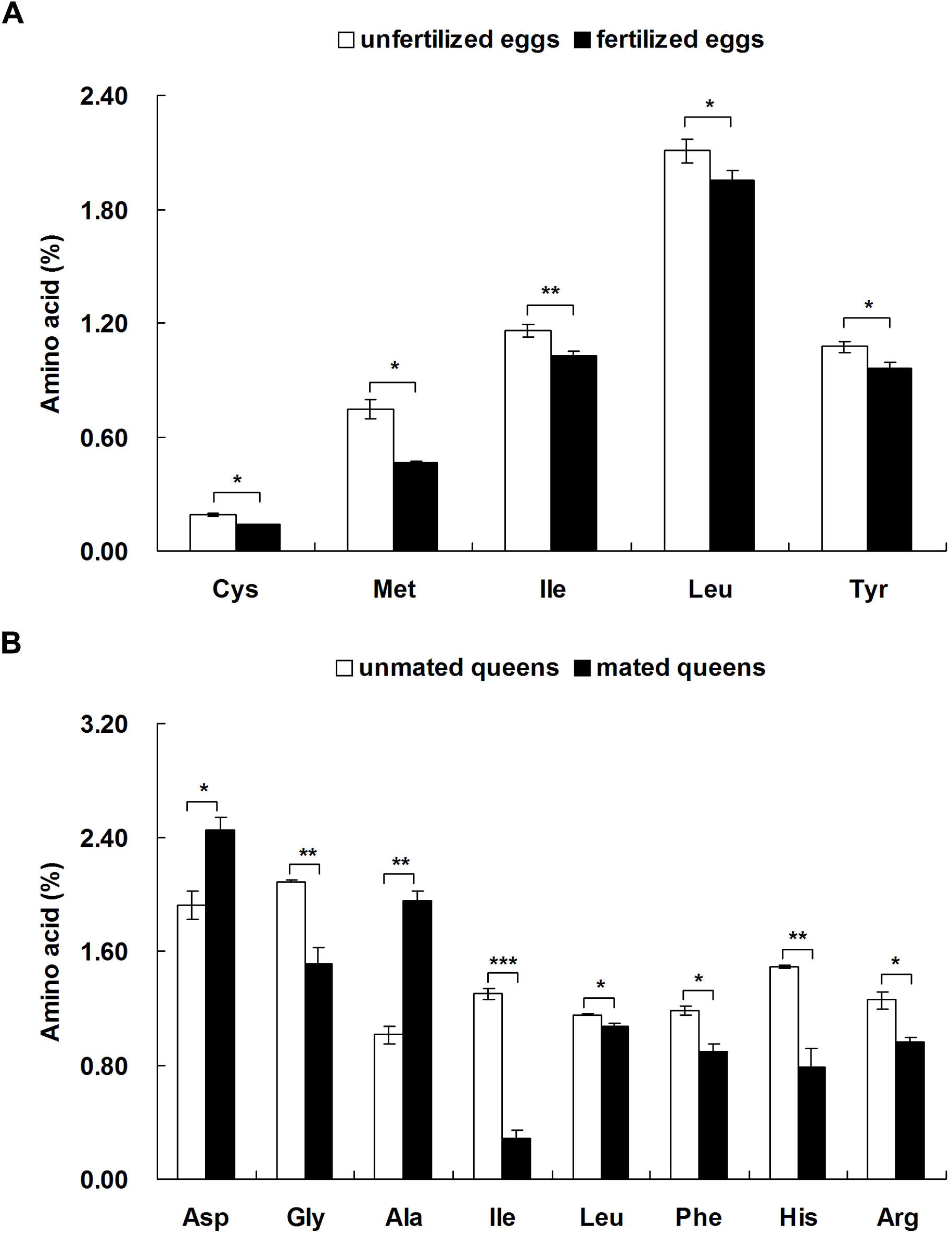
Differences in the levels of 11 amino acids between unfertilized eggs/unmated queens and fertilized eggs/mated queens.

### Trace elements

The Ca level in unfertilized eggs from FF colonies was significantly higher than in fertilized eggs from FM colonies (Fig. 4A; *t* = -4.923, *df* = 2, *p* = 0.039), but the levels of five trace elements (Na, K, Zn, Fe and Mn) in unfertilized eggs were significantly lower than in fertilized eggs (Fig. 4A; Na: *t* = 26.179, *df* = 2, *p* = 0.001; K: *t* = 7.703, *df* = 2, *p* = 0.016; Zn: *t* = 4.436, *df* = 2, *p* = 0.047; Fe: *t* = 5.079, *df* = 2, *p* = 0.037; Mn: *t* = 5.038, *df* = 2, *p* = 0.037). The levels of four trace elements (Zn, Fe, K and Na) in unmated queens were significantly or marginally significantly higher than in mated queens (Fig. 4B; Zn: *t* = 4.080, *df* = 2, *p* = 0.015; Fe: *t* = 3.075, *df* = 2, *p* = 0.037; K: *t* = 3.203, *df* = 2, *p* = 0.033; Na: *t* = 2.163, *df* = 2, *p* = 0.097). However, the Mn level in unmated queens was significantly lower than in mated queens (Fig. 4B; *t* = -3.878, *df* = 2, *p* = 0.018).

**Fig 4:**
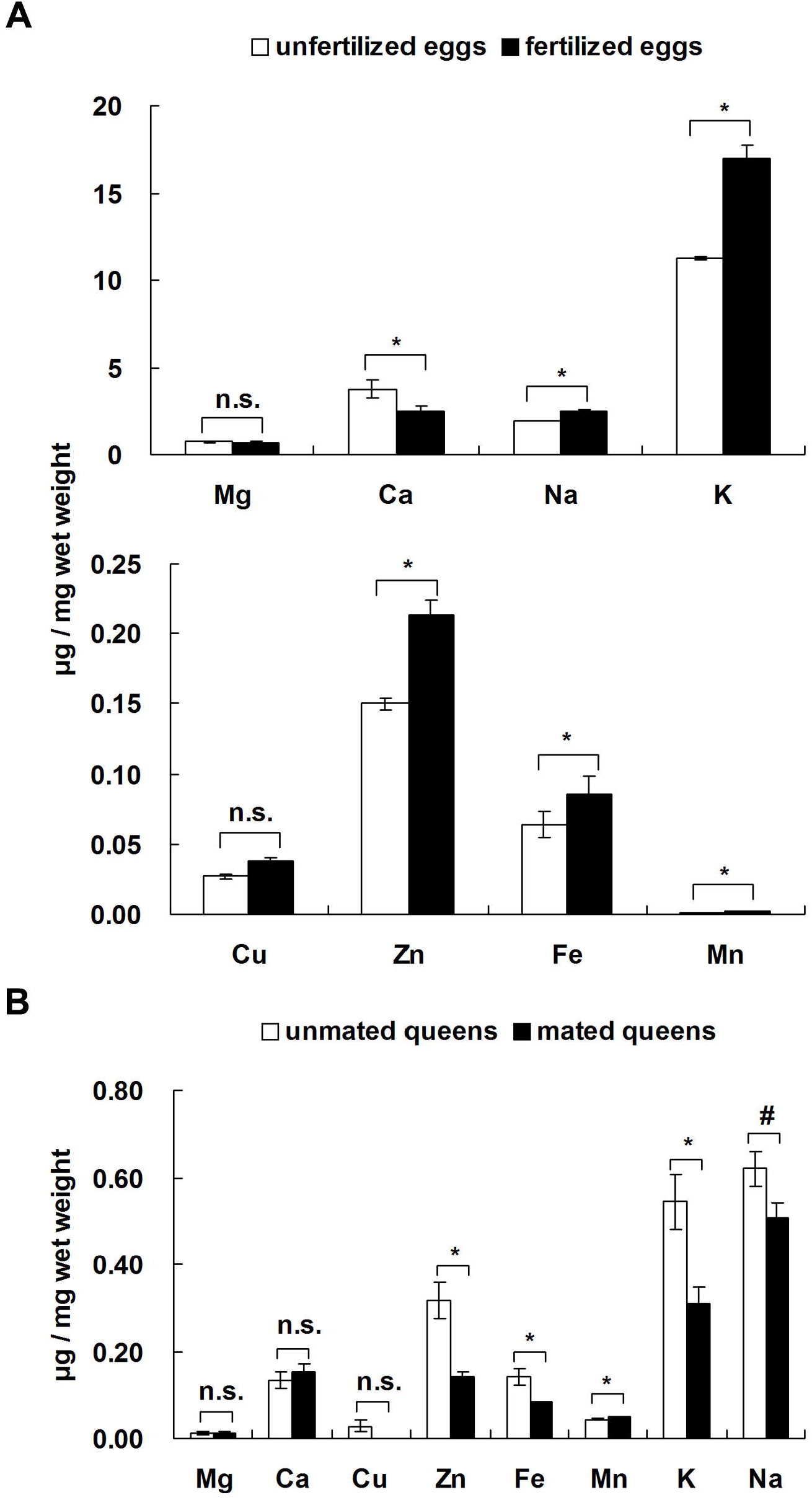
Differences in the levels of eight trace elements between unfertilized eggs/unmated queens and fertilized eggs/mated queens.

### Nutrient content

The levels of proteins and cholesterol in unfertilized eggs from FF colonies were significantly higher than in fertilized eggs from FM colonies (Fig. 5A, protein: *t* = 4.038, *df* = 4, *p* = 0.016; Fig. 5B, cholesterol: *t* = 3.500, *df* = 4, *p* = 0.025), but the glucose level of unfertilized eggs was significantly lower than in fertilized eggs (Fig. 5C; *t* = -6.124, *df* = 4, *p* = 0.004). There were no significant differences in the triglyceride level between unfertilized and fertilized eggs, but the triglyceride level of unmated queens was significantly lower than in mated queens (Fig. 5D; *t* = -2.906, *df* = 4, *p* = 0.044).

**Fig 5:**
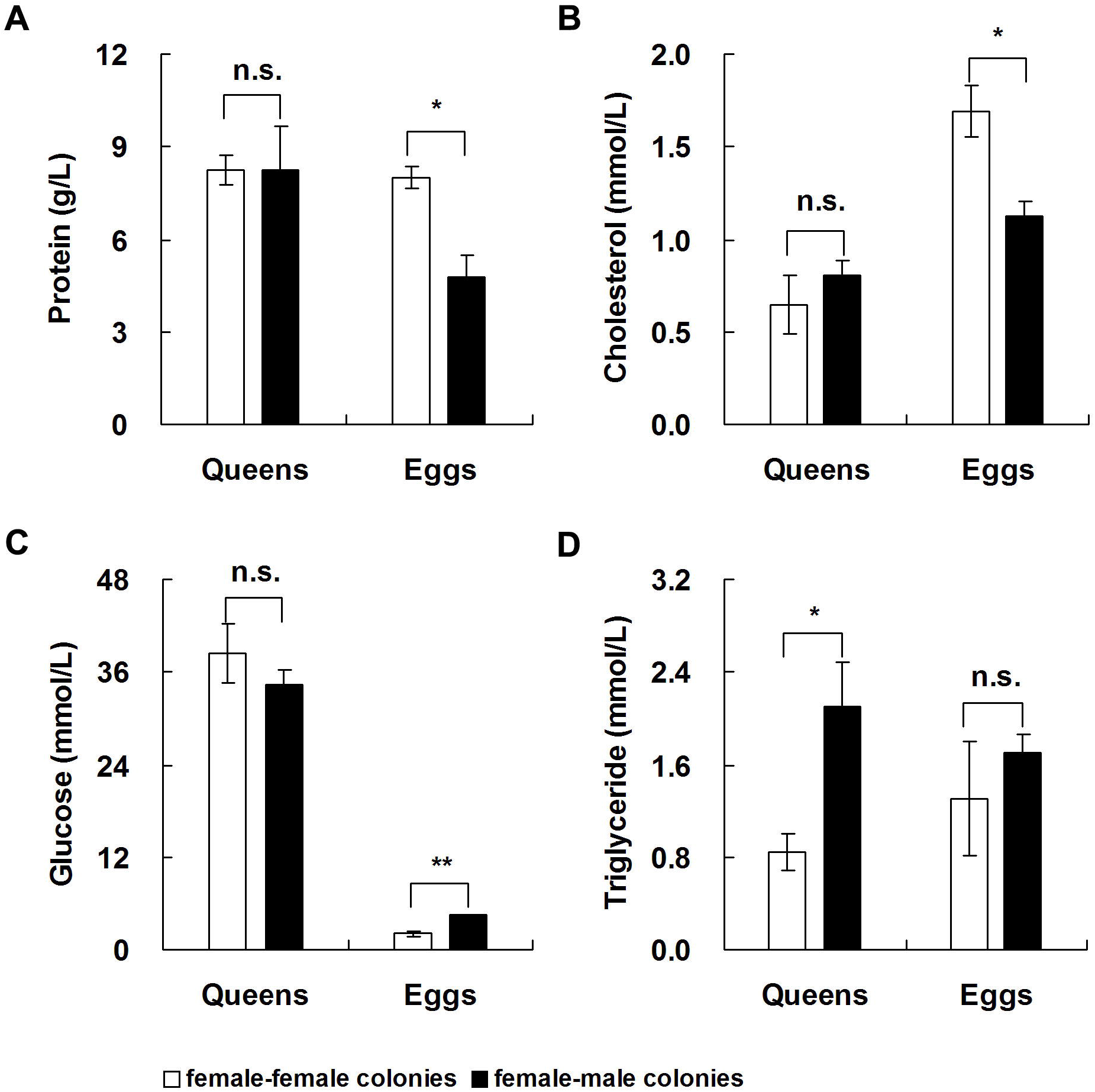
Differences in the content of four nutrients between unfertilized eggs/unmated queens and fertilized eggs/mated queens.

### Hormones and neurotransmitters

Levels of the neurotransmitter serotonin in unmated queens from FF colonies were significantly lower than in mated queens from FM colonies (Fig. 6; *t* = -5.867, *df* = 4, *p* = 0.004), but there were no significant differences in the levels of two hormones (JH III and ecdysone) or other two neurotransmitters (octopamine and dopamine) between unmated and mated queens (Fig. 6).

**Fig 6:**
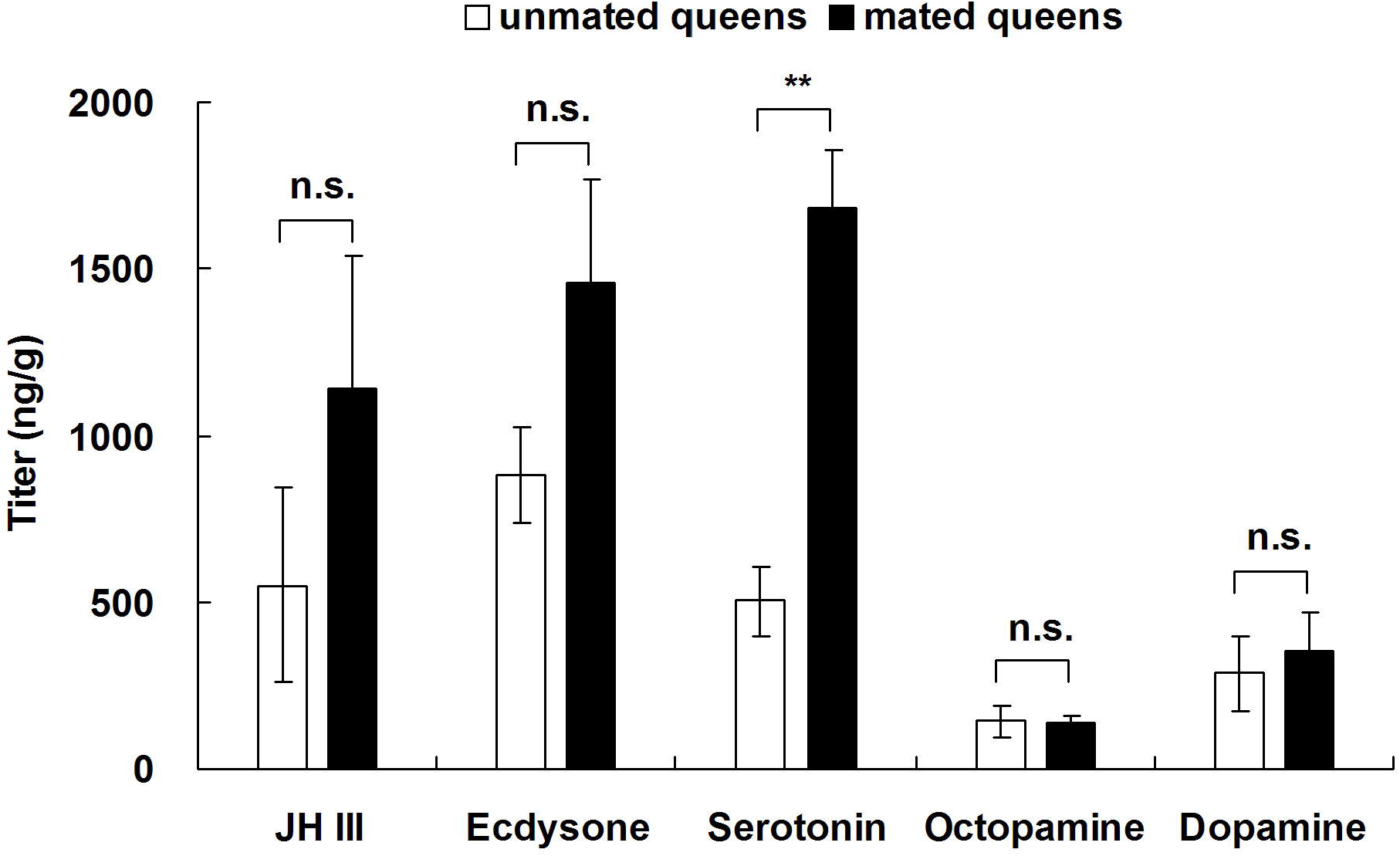
Differences in the levels of two hormones and three neurotransmitters between unmated and mated queens.

### Reproductive genes

The expression of three reproductive genes in unfertilized eggs from FF colonies was significantly higher than in fertilized eggs from FM colonies (Fig. 7A; *vtg 1*: *t* = 6.319, *df* = 2, *p* = 0.024; *rab 11*: *t* = 17.528, *df* = 2, *p* = 0.003; *JHE 1*: *t* = 15.400, *df* = 2, *p* = 0.004). Moreover, the expression levels of two reproductive genes in unmated queens were significantly higher than in mated queens (Fig. 7B; *rab 11*: *t* = 6.614, *df* = 2, *p* = 0.022; *JHE 1*: *t* = 4.510, *df* = 2, *p* = 0.046).

**Fig 7:**
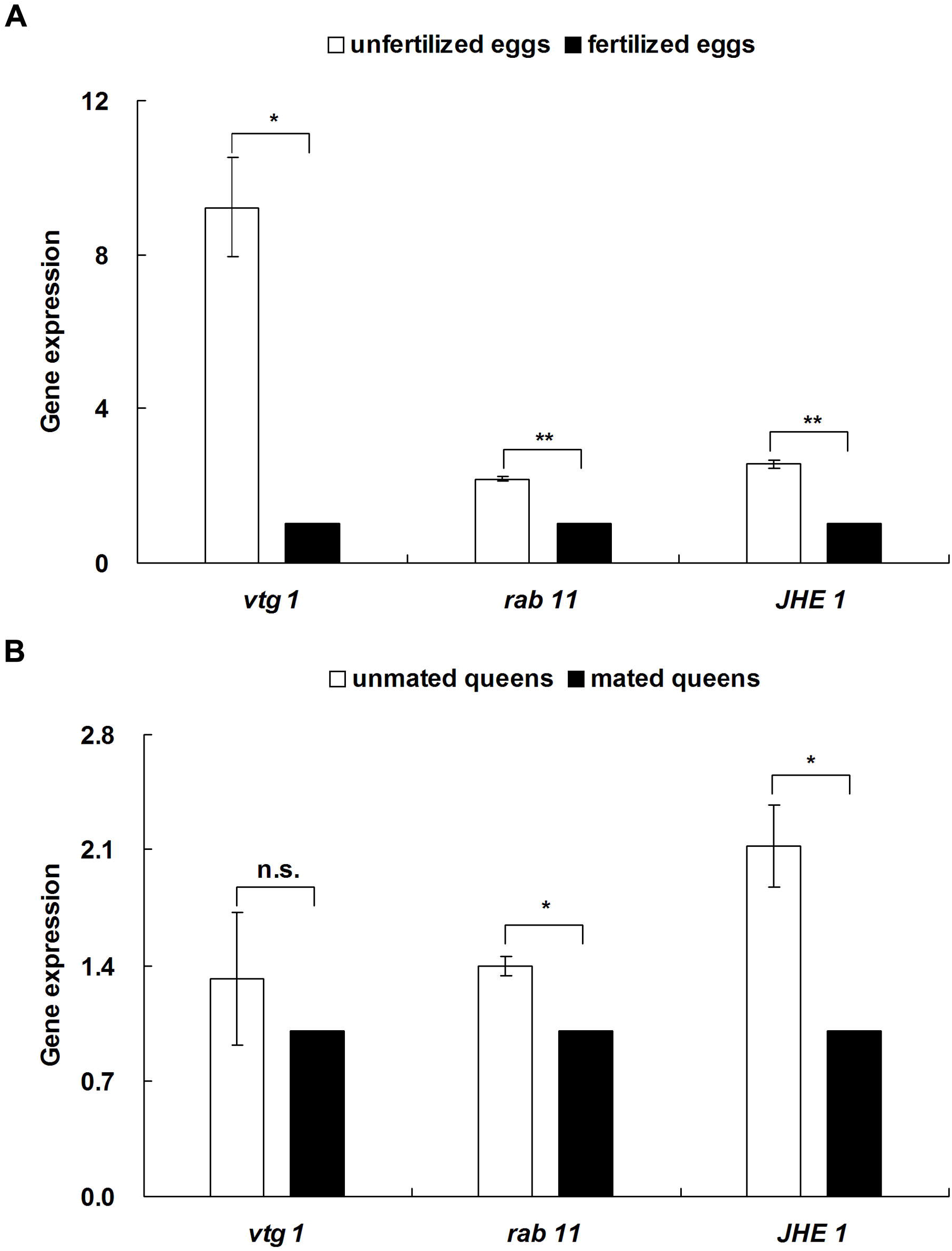
Differences in the expression of three reproductive genes between unfertilized eggs/unmated queens and fertilized eggs/mated queens.

## DISCUSSION

In this study, none of the unfertilized eggs produced by unmated queens from colonies of *R. chinensis* hatched under laboratory or simulated field conditions, suggesting that parthenogenesis does not occur in *R. chinensis*, which is consistent with inferences from the presence of heterozygote genotypes in neotenic reproductives of *R*. *chinensis* (Huang et al., 2013) and the lack of a female-biased sex ratio in the flying alates of *R. chinensis* (Li et al., 2015). Our results indicated that the embryos of all unfertilized eggs ceased growth prior to stage V, potentially due to a lack or reduction of certain important physiological substances. In addition, almost all of the unfertilized eggs (14/15) lacked micropyles but all of the fertilized eggs had micropyles in *R. chinensis*; parthenogenetic eggs of *R. speratus* also lack micropyles (Yashiro and Matsuura, 2014).

Amino acids play an important role in ovarian and embryonic development in animals (Osako et al., 2007; Sato et al., 2009). We found that the levels of five amino acids (Ile, Leu, Cys, Met and Tyr) in unfertilized eggs were significantly higher than in fertilized eggs. A possible explanation for this result is that the sustained embryonic growth of fertilized eggs requires considerable quantities of these five amino acids to complete incubation. These data are consistent with observations in mice that embryonic growth requires substantial levels of Cys (De Matos et al., 2003). Mated queens had significantly lower levels of six amino acids (Gly, Ile, Leu, Phe, His and Arg), suggesting that mated queens must consume large quantities of these amino acids to complete ovarian development after mating in *R. chinensis*. Compared with unmated queens, the significantly higher levels of two amino acids (Asp and Ala) in mated queens may be derived from male reproductives via mating or trophallaxis.

Several ions, including Ca, K, Na, Cu and Zn, are important for growth, ovarian development, fecundity, and embryonic growth in insects (Judd et al., 2010; McFarlane, 1991). The levels of four trace elements (Na, K, Zn and Fe) in fertilized eggs were significantly higher than in unfertilized eggs, but their levels in mated queens were significantly lower than in unmated queens. A possible explanation for this phenomenon is that mated queens transfer their Na, K, Zn and Fe to fertilized eggs. These four trace elements may be necessary during embryonic growth and incubation. These data are consistent with a previous study that found that a decrease in Zn and/or Fe levels resulted in the cessation of growth in human embryos (Bai et al., 2014). The significantly higher levels of Mn in fertilized eggs and mated queens indicated that Mn plays an important role in ovarian and embryonic development during sexual reproduction. These data are consistent with the observation that a lack of Mn decreases the hatchability of eggs in chickens (Leach and Gross, 1983). In addition, the Ca level in fertilized eggs was significantly lower than in unfertilized eggs, suggesting that fertilized eggs require substantial levels of Ca to sustain embryonic growth during incubation. Similarly, normal embryonic development of eggs in turtles, crocodiles and birds consumes considerable quantities of Ca and obtains a portion of the Ca from the eggshell (Dunn and Boone, 1977).

Macronutrients (protein, glycogen, and lipid) play a crucial role in embryonic growth (Diss et al., 1996; Sloggett and Lorenz, 2008) in insects. In the present study, the levels of proteins and cholesterol in fertilized eggs were significantly lower than in unfertilized eggs, suggesting that the sustained embryonic growth and incubation of fertilized eggs requires more protein and cholesterol than in unfertilized eggs (Diss et al., 1996; Moran, 2007). However, the glucose level in fertilized eggs was significantly higher than in unfertilized eggs, implying that glucose may be a primary energy source in the late stages of embryonic growth (Ding et al., 2008). Mated queens had significantly higher levels of triglycerides than unmated queens, suggesting that triglycerides are an important energy source during ovarian development after mating in *R. chinensis*. This result is consistent with the rapid increase in triglycerides observed during ovarian development in the mated female adults of *Marsupenaeus japonicus* (Cavalli et al., 2001). A portion of the triglycerides in mated queens may be derived from males via mating or trophallaxis (Matsuura and Nishida, 2001; Shellman-Reeve, 1990).

Little is known about which genes are involved in the regulation of ovarian and embryonic growth in termites (Ishitani and Maekawa, 2010; Maekawa et al., 2010). In the present study, we found that the expression levels of two reproductive genes (*rab 11* and *JHE 1*) in unfertilized eggs and unmated queens were significantly higher than in fertilized eggs and mated queens. A previous study demonstrated that *Rab 11* is a small GTP binding protein involved in vesicular trafficking and plays a crucial role in the fertility of *Drosophila* (Tiwari et al., 2008). Juvenile hormone esterase (JHE), a member of the carboxylesterase family, contributes to the rapid decline in JH in most of insects. JH is well known to play an important role in the reproductive competence and caste polyphenism of termites (Cornette et al., 2008; Elliott and Stay, 2008; Zhou et al., 2007). Thus, we predicted that these two reproductive genes (*rab 11* and *JHE 1*) might play a role in the ovarian and embryonic development of *R. chinensis*. In this study, we did not find significant differences in two other hormones (JH III and ecdysone) between mated queens and unmated queens, but the serotonin level of mated queens was significantly higher than in unmated queens. As a neurotransmitter, serotonin is correlated with reproductive behaviour in mice (Liu et al., 2011; Zhang et al., 2013), suggesting that it might also play an important role in the sexual reproduction of *R. chinensis*. Vitellogenin is the dominant egg yolk protein in insects (Sappington and Raikhel, 1998). The expression of the gene *vtg 1* in fertilized eggs was significantly lower than in unfertilized eggs in this study, suggesting that fertilized eggs require more vitellogenin to complete embryonic development and incubation.

In conclusion, unfertilized eggs can be produced by unmated queens of *R. chinensis* but do not hatch under laboratory and simulated field conditions, suggesting that *R. chinensis* exhibits neither AQS nor parthenogenesis. Through physiological analyses, we found that unfertilized eggs ceased embryonic growth and had significant differences in morphological characters, size and micropyle number compared with fertilized eggs in the final stage of development. Moreover, fertilized eggs consumed significantly larger quantities of five amino acids (Cys, Met, Ile, Leu and Tyr), Ca, protein, and cholesterol during embryonic growth and incubation. Fertilized eggs may obtain a portion of four trace elements (Na, K, Zn and Fe) from mated queens, which facilitate embryonic growth during incubation. The significantly higher levels of Mn and glucose in fertilized eggs than unfertilized eggs suggest that these substances play an important role in the completion of embryonic growth and incubation. The significantly higher levels of triglycerides and serotonin in mated queens than in unmated queens imply that they are very important for ovarian development during sexual reproduction. The greater expression of three genes (*vtg1*, *rab11* and *JHE1*) in unfertilized eggs and two genes (*rab11* and *JHE1*) in unmated queens suggests that these reproductive genes may be involved in the regulation of ovarian and embryonic growth in *R. chinensis*. Overall, the differences in these physiological indices may substantially affect ovarian and embryonic growth and prohibit the incubation of unfertilized eggs in *R. chinensis*. Our physiological findings contribute to an understanding of why the subterranean termite *R. chinensis* exhibits neither AQS nor parthenogenesis even though it is a close relative of *R. speratus*, which exhibits AQS.

## MATERIALS AND METHODS

### Formation of female-female colonies and female-male colonies under laboratory conditions

The alates of *R. chinensis* used in this study were collected together with nest wood just prior to the swarming season. In April 2013, four mature colonies were collected from Nanwang Hill (A), Yujia Hill (B), and Shizi Hill (C and D), in Wuhan City, China. The termites were reared in an open plastic container (670×480×410 mm^3^) covered by nylon mesh in a dark room with a temperature of 25°C for 14 days and then moved to a room with a temperature of 30°C to help the alates fly (Matsuura et al., 2002; Matsuura and Nishida, 2001). Details of the methods for founder combinations and brood observation are described in Text S1.

### Formation of female-female colonies and female-male colonies under simulated field conditions

In April 2012, three mature colonies were collected from Nanwang Hill (E), Yujia Hill (F), and Shizi Hill (G), in Wuhan City, China. We used the same method as described above for the laboratory conditions to help alates fly. Details of the methods for founder combinations and brood observation are described in Text S1.

### Morphological observation of eggs at the five developmental stages

We used the same colonies and method as described above for the laboratory conditions to build founder combinations. Details of the methods for egg culture and observation of embryonic growth, egg sizes and micropyle number are described in Text S1.

### Amino acids

Unmated queens and unfertilized eggs from FF colonies and mated queens and fertilized eggs from FM colonies were sampled for the analysis of amino acid composition using an automatic amino acid analyser. Details of the methods for quantifying amino acids are described in Text S1.

### Trace elements

Unmated queens and unfertilized eggs from FF colonies and mated queens and fertilized eggs from FM colonies were sampled for the analysis of levels of eight trace elements—Ca, Cu, Fe, K, Mg, Mn, Na, and Zn—using ICP-OES (Judd et al., 2010). Details of the methods for the assays of trace elements are described in Text S1.

### Nutrient content

Unmated queens and unfertilized eggs from FF colonies and mated queens and fertilized eggs from FM colonies were sampled for the analysis of levels of four nutrients—protein, cholesterol, glucose and triglycerides—using a microplate spectrophotometer (Li et al., 2015). Details of the methods for nutrient content analysis are described in Text S1.

### Hormones and neurotransmitters

Unmated queens from FF colonies and mated queens from FM colonies were sampled to determine the levels of hormones (JH III and ecdysone) and neurotransmitters (serotonin, octopamine and dopamine) in unmated and mated queens using multiple reaction monitoring (MRM) analysis. Details of the MRM analysis are described in Text S1.

### Reproductive genes

Unmated queens and unfertilized eggs from FF colonies and mated queens and fertilized eggs from FM colonies were sampled to assess the expression of three reproductive genes—*Vitellogenin 1* (*vtg 1*), *rab 11* and juvenile hormone esterase-like protein Est1 (*JHE 1*)—using qPCR. Details of the methods for RNA extraction, primer design, PCR application and qPCR are described in Text S1.

## Acknowledgements

We thank Fei Chu, Wei Wang, Huan Xu and Yongyong Gao for their assistance with field collections. We thank the anonymous reviewers for providing valuable comments on earlier drafts of this manuscript.

## Author contributions

G.H.L., C.L.L. and Q.Y.H. conceived and designed the experiments; G.H.L., L.L., P.D.S. and Q.Y.H. performed the experiments; G.H.L., L.L. and Q.Y.H. analysed the data; G.H.L., L.L., P.D.S., Y.W., X.W.C and Q.Y.H. wrote the paper. All authors read and approved the final manuscript.

